# The Decision Decoding ToolBOX (DDTBOX) – A multivariate pattern analysis toolbox for event-related potentials

**DOI:** 10.1101/153189

**Authors:** Stefan Bode, Daniel Feuerriegel, Daniel Bennett, Phillip M. Alday

**Affiliations:** Melbourne School of Psychological Sciences, The University of Melbourne, Australia; School of Psychology, Social Work and Social Policy, University of South Australia, Australia; Princeton Neuroscience Institute, Princeton University, Princeton, New Jersey, USA; Department of Psychology of Language, Max Planck Institute for Psycholinguistics, Nijmegen, The Netherlands

**Keywords:** multivariate pattern classification analysis (MVPA), decoding, event-related potentials (ERPs), electroencephalography (EEG), toolbox, support vector machines

## Abstract

In recent years, neuroimaging research in cognitive neuroscience has increasingly used multivariate pattern analysis (MVPA) to investigate higher cognitive functions. Here we present DDTBOX, an open-source MVPA toolbox for electroencephalography (EEG) data. DDTBOX runs under MATLAB and is well integrated with the EEGLAB/ERPLAB and Fieldtrip toolboxes (Delorme and Makeig, 2004; Lopez-Calderon and Luck, 2014; Oostenveld et al. 2011). It trains support vector machines (SVMs) on patterns of event-related potential (ERP) amplitude data, following or preceding an event of interest, for classification or regression of experimental variables. These amplitude patterns can be extracted across space/electrodes (spatial decoding), time (temporal decoding), or both (spatiotemporal decoding). DDTBOX can also extract SVM feature weights, generate empirical chance distributions based on shuffled-labels decoding for group-level statistical testing, provide estimates of the prevalence of decodable information in the population, and perform a variety of corrections for multiple comparisons. It also includes plotting functions for single subject and group results. DDTBOX complements conventional analyses of ERP components, as subtle multivariate patterns can be detected that would be overlooked in standard analyses. It further allows for a more explorative search for information when no ERP component is known to be specifically linked to a cognitive process of interest. In summary, DDTBOX is an easy-to-use and open-source toolbox that allows for characterising the time-course of information related to various perceptual and cognitive processes. It can be applied to data from a large number of experimental paradigms and could therefore be a valuable tool for the neuroimaging community.

## 1. General introduction

In recent years, the use of multivariate pattern analysis (MVPA) techniques for neuroimaging data has rapidly increased. Beginning with Edelman and colleagues’ (1998) and Haxby and colleagues’ (2001) seminal studies, applications of MVPA to functional magnetic resonance imaging (fMRI) data have become increasingly popular, leading to a strong trend towards investigating representations and predicting “information” (i.e. the content of cognition) in cognitive neuroscience (for reviews and comments see: Davis and Poldrack, 2013; Haynes, 2015; Haynes and Rees, 2006; Heinzle et al., 2012; Hogendoorn, 2015; Mur et al., 2009; Norman et al., 2006; Kriegeskorte et al., 2006; Tong and Pratte, 2012; Woolgar et al., 2014; 2016) and to the publication of several toolboxes (Hanke et al., 2009; Hebart et al., 2015; Oosterhof et al., 2016). Most variants of MVPA have in common that (usually supervised) classifiers are trained to predict the content of cognitive processes directly from local spatial activation patterns, as measured by the blood-oxygen-level-dependent (BOLD) signal. Due to the poor temporal resolution of fMRI, however, it is often difficult to track any fast decision processes in time, and to determine the timecourse with which predictive information regarding these processes is represented in neural data.

One potential solution to this problem is to apply MVPA to electroencephalography (EEG) data, as EEG has a far better temporal resolution in the range of milliseconds, as opposed to seconds in fMRI (Other techniques such as magnetoencephalography, MEG, will not be discussed here; for reviews see King and Dehaene, 2014; Grootswagers et al., 2016). Multivariate analysis techniques were already applied to EEG data to investigate cognition several decades ago (Gevins et al., 1979), but are currently experiencing a strong revival (for reviews and technical comments see: Bai et al., 2007; Blankertz et al., 2011; Contini et al., in press; King and Dehaene, 2014; Parra et al., 2005; Sajda et al., 2009; Stokes et al., 2015). By taking advantage of the multivariate nature of EEG signals, multivariate analysis techniques can, for example, predict the outcomes of decisions, parameters of decision models, and decision errors directly from activity patterns (e.g., Blank et al., 2013; Bode et al., 2012; 2014; Bode and Stahl, 2014; Boldt and Yeung, 2015; Charles et al., 2014; Chung et al., 2015; Das et al., 2010; Parra et al., 2002; Philiastides and Sadja, 2006; Philiastides et al., 2006; Ratcliff et al., 2009; Tzovara et al., 2015; van Vugt et al., 2012; for related approaches see: El Zein et al., 2015; Wyart et al., 2012; 2015). Others have used similar techniques to investigate visual awareness (Hogendoorn et al., 2015; Hogendoorn and Verstraten, 2013; Fahrenfort et al., 2017), multisensory integration (Chan et al., 2017), and automatic processing of semantic features of task-irrelevant stimuli (Bode et al., 2014). MVPA has also been extensively used for constructing brain-computer interfaces (BCIs; e.g., Müller et al., 2004, 2008).

We present a novel open-source toolbox for MATLAB––the Decision Decoding ToolBOX (DDTBOX)—that performs MVPA on the high-temporal-resolution EEG data typically analysed using univariate analyses of event-related potentials (ERPs). However, instead of analysing signals at single electrodes (i.e. channels), or averaging across a group of selected electrodes, for which ERP components have been described and linked to specific cognitive processes (e.g., Luck, 2005), MVPA is applied to data from all electrodes in a predefined analysis time window, which thereby serves as multivariate input to a classifier (Parra et al., 2005). Such patterns of amplitude data can be extracted across space (e.g., from data averaged over a time window for each channel: spatial decoding), time (e.g., from all timepoints for each channel separately: temporal decoding), or both (from all timepoints for all channels: spatiotemporal decoding). This approach is arguably often more data-rather than hypothesis-driven compared to conventional ERP analyses, and has several benefits: First, even subtle multivariate EEG patterns that differ between experimental conditions can be detected that would otherwise be overlooked (Bode et al., 2012). Second, single-trial patterns of activity can be directly linked to parameters in decision models (e.g., Philiastides and Sadja, 2006; Ratcliff et al., 2009; Tzovara et al., 2015), or used to predict subjective properties of stimuli such as arousal (e.g., Bode et al., 2014). Finally, MVPA on ERPs allows for a more explorative search for information when no ERP component is known to be specifically linked to the cognitive process of interest; it does not require *a priori* knowledge of the location and timing of an effect, which can vary substantially across experiments (discussed in Groppe et al., 2011).

We will now introduce the functionality of DDTBOX, which can be applied to data from a variety of experimental paradigms (and is by no means restricted to decision-making research). DDTBOX requires minimal experience with MATLAB coding, and integrates well with EEGLAB/ERPLAB (Delorme and Makeig, 2004; Lopez-Calderon and Luck, 2014) and FieldTrip (Oostenveld et al. 2011), allowing users to prepare data using standard preprocessing pipelines for ERP analyses with only minimal additions. DDTBOX can, however, also use data preprocessed with many other commercially available software packages. In the following, we will first briefly discuss basic principles of classification approaches for the study of EEG signals. Then, we will provide an overview of the general architecture of the DDTBOX, complementing our detailed online user manual <u>(https://github.com/DDTBOX/DDTBOX/wiki).</u> This will be followed by a user-oriented introduction to using DDTBOX, covering the general principles and features, the functional structure of the toolbox, and a section on how to prepare EEG data and how to configure analyses in DDTBOX. We will then provide examples of research which has already used beta-versions of DDTBOX, as well as limitations of analyses offered by DDTBOX. Finally, we will conclude by giving an outlook into planned future developments and extensions, as well as options for users to directly contribute to the toolbox. Where relevant, we will make reference to the detailed, more technical documentation available online, which is designed as an additional hands-on guide to the DDTBOX. We have also provided an example dataset online that can be downloaded to learn how to use DDTBOX, available at <u>https://osf.io/bndjg/.</u>

## 2. Classification based on ERP data

Machine learning has recently gained strong popularity in systems neuroscience. In particular, a supervised-learning approach using support vector machines (SVMs) has proved to be a powerful tool for neuroimaging analysis (e.g., Haynes, 2015; Grootswagers et al., 2016). The power of this approach is derived from the fact that in most experiments, experimenters know the categories of interest *a priori*. These categories of interest typically correspond to different experimental conditions or participant response types (e.g., participants make decisions regarding object categories, make errors versus correct responses, report different subjective experiences, etc.). The aim for analysis is then to find patterns of neural activity that distinguish these categories. For multivariate EEG signals, this would correspond to finding patterns in the signal across time and space (electrodes) that distinguish the categories of interest. While this approach is still correlative, in the sense that it seeks to identify patterns of covariance between neural data and latent cognitive variables, its great advantage is that the structure of the neural data need not map straightforwardly to known aspects of the cognitive variables. Instead, it is sufficient that the EEG signal patterns *predict* the cognitive variables, thereby permitting researchers to conclude that information regarding cognitive variables is present in the neural data, either decodable from specific electrodes, or from specific processing time windows (Yarkoni et al., in press).

A detailed description of SVMs has been provided elsewhere (Cortes and Vapnik, 1995; Burges, 1998; Hastie, Tibshirani and Friedman, 2001). Put simply, the general principle of SVM classification is to construct a hyperplane (i.e. a decision boundary) in multidimensional feature space to optimally separate exemplars into different categories (i.e. neural data mapping onto different experimental conditions). The further the separating hyperplane from the nearest elements of each group (known as the “support vectors”), the better the classification. To avoid circularity (Kriegeskorte et al. 2009), estimation of the hyperplane must be performed on data independent from test data (left-out data from the same experiment, or new data from an identical experiment), which is subsequently used to evaluate the model by assigning category “labels” to each exemplar in the test data set. This usually involves k-fold cross-validation, in which the data are divided into *k* subsets. The classifier is trained on k-1 subsets and tested using the left-out subset. This procedure is then independently repeated with each subset serving as the test data set once while training on the others (for an example see Meyers and Kreiman, 2011). The average accuracy across all cross-validation steps, often referred to as “classification accuracy” or “decoding accuracy”, can then be treated as an index for whether information about the categories of interest was represented in this specific pattern of brain activity. Statistically, this question can be assessed by submitting classification accuracy values to statistical testing, either against a theoretical chance performance level (e.g., with two balanced classes, the expected chance level = 50%), or against an empirical test distribution, e.g., by comparing against results of analyses using randomly-shuffled condition labels (e.g., Stelzer et al., 2013). Although SVM classification is binary in nature, it can easily be extended to more complex multi-class classification problems by combining results from all pair-wise class combinations or performing one-vs.-other comparisons.

DDTBOX can also perform a type of generalisation analysis, testing whether patterns of information that discriminate between categories are stable across experimental contexts. For this, the classifiers are trained on data from one context (e.g., correct and error responses in task A) and used to predict the classes from data from another context (e.g., correct and error responses in task B). Such an approach is known as a cross-condition classification analysis. This can reveal whether the neural patterns that discriminate between outcomes are consistent across different task or stimulus presentation conditions, as the classifier should only be able to generalise from one to the other is patterns are highly similar (for an example from fMRI, see Bode et al., 2013).

For cognitive variables of interest that are continuous rather than categorical (such as response times), an alternative to SVM classification is support vector regression (SVR). SVR allows for trial-by-trial values of a continuous variable to be mapped to predicted values of that variable. DDTBOX offers both options, as we will outline below.

## 3. General principles of DDTBOX

In order to perform a classification analysis, DDTBOX first requires the user to define discrimination groups, corresponding to categories of interest. These could be experimental conditions (e.g., different object categories) but also participants’ behaviour (e.g., correct and incorrect responses). The event-locked ERP data, which is used for the analysis, has undergone all pre-processing steps such as artefact correction and is epoched into time periods of interest as for conventional ERP analyses. Events can be either exogenous, such as stimulus presentations, or endogenous, such as behavioural responses. The epochs of ERP data are then sorted with respect to the categories of interest, and each epoch is assigned a class label corresponding to its category.

### 3.1 SVM back-end software, types of analysis, and kernels

DDTBOX’s main function is to prepare exemplars of patterns of ERP data for each participant for SVM classification or regression analyses. To perform such analyses, DDTBOX interacts with existing machine learning software packages to perform classification/regression (similar to The Decoding Toolbox for fMRI; Hebart et al., 2015). The user can choose between the LIBSVM package (Chang and Lin, 2011), which has been used by other toolboxes in the field (e.g., Hebart et al., 2015), and LIBLINEAR (Fan et al., 2008), a less flexible but faster implementation of commonly-used SVM classification and regression algorithms (see the online documentation for details). Several SVM analysis options are available, including different SVM fitting methods and kernels. We refer to the websites of these software packages for detailed explanations of these options (LIBSVM: <u>https://www.csie.ntu.edu.tw/~cjlin/libsvm;</u> LIBLINEAR: <u>https://www.csie.ntu.edu.tw/~cjlin/liblinear).</u> For most research questions requiring classification, C-SVM (as implemented in LIBSVM) with a linear kernel and a default regularising parameter C = 1 appears to be adequate and standard in the field, and is therefore the default option in DDTBOX. For multivariate regression, DDTBOX uses SVR in LIBSVM with a linear kernel and regularisation parameter C = 0.1 as the default option.

### 3.2 Analysis time window width

DDTBOX performs analyses using a moving window approach: the signals of interest during a prespecified analysis time window are extracted and analysed, and the analysis time window is then moved by a specified step-size through the epoch (depicted in Figure 1A). The user can specify the analysis window width and step-size. The optimal analysis time window width depends on the research question of interest, as information relating to some cognitive processes might be better captured by longer analysis time windows, while other short duration cognitive processes might be better captured using short analysis time windows. Our own previous work has successfully utilised analysis time windows ranging from 10ms (e.g., Bode and Stahl., 2014) to as long as 80ms (e.g., Bode et al., 2012).

**Figure 1.**
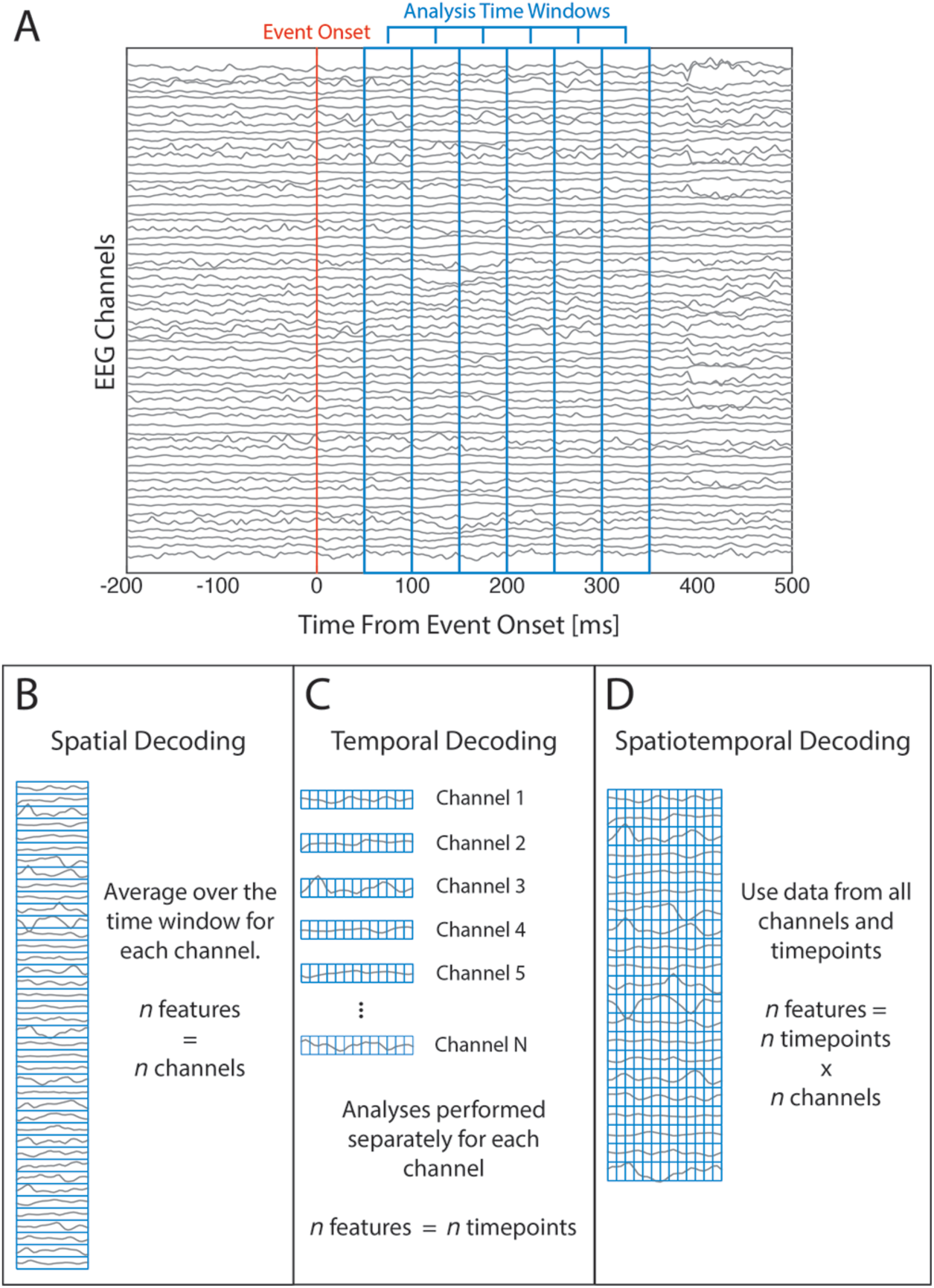
Decoding approaches in DDTBOX. A) Example of the windowed analysis approach. DDTBOX performs MVPA on time windows of EEG data (time windows outlined in blue). For each analysis the time window is moved through the trial by a predefined step size. B) Example of spatial decoding. For each channel EEG data is averaged across timepoints within the analysis time window, resulting in one value per channel used for MVPA. C) Example of temporal decoding. MVPA is performed using data from each timepoint within the analysis time window, for each channel separately. D) Example of spatiotemporal decoding. All timepoints at all channels are used in combination for MVPA.

### 3.3 Analysis time window step size

The analysis time window is moved through the trial at a user-defined step size, independently repeating decoding analyses each time with data from the new time window (depicted in Figure 1A). The step size could be the same as the analysis window width to achieve non-overlapping analysis time windows (e.g., 10ms windows moved in steps of 10ms). Alternatively, the step size could be finer than the window width (e.g., 20ms windows moved in steps of 10ms), leading to partial overlap of analysis time windows. This can be useful, for example, when one is interested in relatively fast cognitive processes, which might occur with a finer temporal resolution than the window size and therefore be captured only partly by two consecutive larger analysis time windows.

### 3.4 Spatial and/or temporal analyses

DDTBOX users can elect to run *spatial analyses* (Figure 1B), which involve averaging across all data points included in the chosen analysis time window for each channel. This procedure results in one data point per channel (*number of channels* x 1 activity pattern). Alternatively, the user can choose to disregard spatial patterns and perform *temporal analyses* (Figure 1C) using data from single channels. In this case, all other channels are ignored, and the data points for the selected channel that are included in the analysis time window (*number of data points x 1*) are treated as the activity pattern of interest. This analysis does not investigate spatially distributed information, but instead focuses on information distributed in time for a given channel. This approach is complementary to spatial classification, but it does not make use of all available (spatial) information. Finally, one can consider both spatial information (over channels) and temporal information (over timepoints) within a chosen analysis time window as the activity pattern (spatiotemporal analyses; *number of data points x number of channels*), as shown in Figure 1D.

### 3.5 Averaging

DDTBOX further provides the user with the option to average across separate sets of exemplars first before training the classifier. The standard option is not to average, which means that usually each experimental trial (or a part of such) is treated as one exemplar for one of the classes of interest. This usually has the advantage of maintaining a large number of exemplars for training and testing. However, if data from a large number of trials are available, one might consider averaging across subsets of trials for the same reasons that averaging is performed to obtain grand average ERPs: to optimise the signal-to-noise ratio. For example, if the experiment was split into 10 separate blocks, one could use block-averaged data for each class instead of single trials (e.g., see Bode et al., 2012). This is similar to first obtaining beta-estimates, or ‘regressors’, for separate functional ‘runs’ in fMRI, and then performing MVPA on these estimates (representing the run-averaged model fit of a general linear model) instead of on single volumes from all trials. Averaging usually results in estimates of exemplars with a higher signal-to-noise ratio, and can improve classification performance in some cases (see Isik et al., 2014; Grootswagers et al., 2016).

### 3.6 Feature weight analyses

DDTBOX allows users to extract and analyse feature weights from the fitted SVM classifiers. Much as regression coefficients describe the contribution of each predictor to the dependent variable, feature weights in SVM describe the contribution of each feature in determining the decision boundary, i.e. separating classes. As such, feature weights are used in DDTBOX to estimate the relative importance of different features (e.g., channels in spatial decoding analyses) for classification or regression. Accordingly, feature weights are analysed in DDTBOX to identify sources of information that the classifier uses to distinguish between experimental categories of interest. The ‘raw’ feature weights derived from SVMs are prone to erroneous interpretations regarding the sources of information used for decoding, as they can be affected by other statistically independent signals (such as noise generated by muscle activity, which as a feature may be strongly weighted but irrelevant). However, this can be corrected in DDTBOX by employing the algorithm described by Haufe and colleagues (2014).

In *spatiotemporal analyses* (see above) the features are single timepoints within the analysis time window for each channel. In DDTBOX feature weights are averaged across timepoints within each analysis window to output an averaged feature weight value for each channel (in consequence, group level feature weight analyses are only implemented for *spatial* and *spatiotemporal analyses*). Furthermore, as the sign of the feature weights indicate the importance of each feature for one or the other (arbitrary) category, and since the sign of each feature weight is therefore arbitrary, DDTBOX computes absolute feature weights, which indicate the importance for the classification in general (i.e. for either category). However, the advanced user can also access the original signed feature weights at individual timepoints within each analysis window. Lastly, feature weights from each analysis step are z-standardised to make them comparable between analyses. Hence, the final output is one absolute, z-standardised feature weight value for each channel for each analysis time window. These are used for group-level statistical testing (see below).

### 3.7 Statistical testing

The result of each single analysis for each participant is a percentage value of correct classifications for all exemplars contained in the test-data set (for classification analyses), or a Fisher-Z transformed correlation between the predicted labels and the true labels (for regression analyses). Then, after the k-fold cross-validation procedure, all k outcome values are averaged to index the overall accuracy. As it is theoretically possible that accuracy estimates were inflated by chance due to the random assignments of exemplars to training and test sets, the default option in DDTBOX is to re-compute the sets m times (i.e. a new, fully independent draw of k sets) and to repeat all analyses for a user-specified number of iterations. The default is to repeat all cross-validated analyses with independently drawn sets ten times. For example, choosing k = 10 for cross-validation, and m = 10 iterations will result in 10 x 10 = 100 analyses, and the final accuracy will be the average of all 100 analyses. This procedure is designed to optimise reliability of results rather than accuracy values.

Statistical testing at a group level is then performed on average accuracy values obtained from the same analysis time window across participants. DDTBOX offers the option of testing against theoretical chance level (e.g., 50% for a balanced two-class classification, 33% for balanced three-class classification, etc.). However, this approach has been criticised recently (Combrisson and Jerbi, 2015). For example, increases in sample variance of accuracy values will also increase the chance of rejecting the null hypothesis when testing against theoretical chance (Allefeld et al., 2016). The default option in DDTBOX is therefore to estimate the empirical chance distribution by running decoding analyses on data with permuted condition labels. Specifically, DDTBOX repeats all original analyses (e.g., m iterations of a k-fold cross-validation procedure) with exactly the same data and the same category labels, but with assignment of labels to exemplars independently randomised for each iteration. This means that any potential biases in the original data (such as unbalanced numbers of exemplars across categories) also affect the permuted-label analyses. The original and the permuted-label analyses are otherwise completely identical, and the results of the permuted-label analyses can then be statistically compared to the original results.

Finally, group decoding accuracy at each analysis time window can be tested for statistical significance using either paired-samples *t*-tests or a group-level analysis method described in Allefeld et al. (2016) based on the minimum statistic (Friston et al., 1999). Importantly, both testing approaches do not provide population inference as do *t*-tests on univariate measures, but instead test the null hypothesis that there are at least some individuals within the sample that show above-chance decoding (i.e. is a fixed-effect analysis; discussed in Allefeld et al., 2016). However, the method based on the minimum statistic also provides lower bound estimates of the prevalence of decodable information in the population.

Analyses run in DDTBOX typically involve a large number of individual tests, requiring corrections for multiple comparisons to control the family-wise error rate. The number of tests performed depends on the number of analysis time windows, which can be minimised by selecting a restricted search space prior to running decoding analyses. DDTBOX offers a variety of correction techniques for multiple comparisons, some of which exploit temporal autocorrelation of the classification accuracy results across time windows to preserve statistical power. Available corrections include the Holm-Bonferroni method (Holm, 1979), maximum statistic and cluster-based permutation tests (Blair and Karniski, 1993; Maris and Oostenveld, 2007), generalised family-wise error rate control (Korn et al., 2004) and false discovery rate control (e.g. Benjamini and Hochberg, 1995; Benjamini et al., 2006). In addition, the distributional assumptions for paired-samples *t-*tests are often violated for samples of classification accuracy scores (Stelzer et al., 2013). DDTBOX can therefore also perform analyses using Yuen’s paired-samples *t-*test (Yuen, 1974; Wilcox, 2012), which is more robust against violations of normality.

DDTBOX further offers group-level statistical testing of feature weights using paired-samples *t-*tests, with corrections for multiple comparisons over channels. Feature weights can be averaged over a number of analysis time windows before statistical testing, if required.

### 3.8 Display options

DDTBOX allows plotting of the decoding performance and feature weight results at various stages. First, users can plot decoding accuracy scores (averaged over cross-validation steps and independent analyses) for individual subjects, for all analysis time windows (*spatial* and *spatiotemporal analyses*) or for all channels within a single time window (*temporal analyses)*. For *spatial* and *spatiotemporal analyses* this is an ‘information time-course’, displaying the average accuracies (y-axis) for each chosen analysis time window (x-axis). Results of permuted condition labels analyses can also be plotted. This could be useful to quickly visually inspect the results for appropriateness of the chosen parameters (such as the window widths or step size), and also to confirm that the shuffled-label control analysis produces chance results. By contrast, *temporal analysis* results are plotted as a spatial map of accuracies for each channel, which are plotted as a heat map with a topographic projection onto the scalp.

Similarly, at a group level information time-course plots can be generated for *spatial* and *spatiotemporal analyses*, displaying the group-level accuracies (and optionally the permuted labels analysis results in the same plot) with error bars denoting standard errors of the mean. Users also have the option to include a vertical bar indicating the timing of the event of interest, as well as automatic marking of statistically significant analysis time windows based on a user-specified alpha level. Axis labels are automatically generated (based on the included baseline period and sampling rate, as well as minimum and maximum accuracy values) but can be manually modified, if desired. The *temporal analyses* group results are again heat maps displaying the colour-coded average group-level accuracy for each channel (note that standard errors are not included in this plot).

For the display of group-level feature weight maps (*spatial* and *spatiotemporal analyses*), two options are available. Firstly, a matrix of z-standardised, absolute feature weights per channel (y-axis) can be displayed for user-selected analysis time windows (x-axis). Secondly, the z-standardised, absolute group-level feature weights can be displayed for single analysis time windows or averages of user-specified analysis time windows. Feature weights can also be plotted as maps thresholded by statistical significance. All figures are plotted using MATLAB plotting routines, can be manually modified if desired, and exported to file formats including TIFF, JPG, PDF, EPS, and many others.

## 4. Functional structure of DDTBOX

The functional structure of DDTBOX is extensively described in the wiki <u>(https://github.com/DDTBOX/DDTBOX/wiki/DDTBOX-Code-Structure)</u> and will not be repeated here in detail. The order of data processing steps in DDTBOX for MVPA on single subject datasets is displayed in Figure 2A. The operations performed in DDTBOX for group-level statistical testing are illustrated in Figure 2B. Advanced users, who might want to gain access to data after specific processing steps, or who are considering expanding the toolbox at specific stages according to their needs, can use this information to easily navigate through the code.

**Figure 2.**
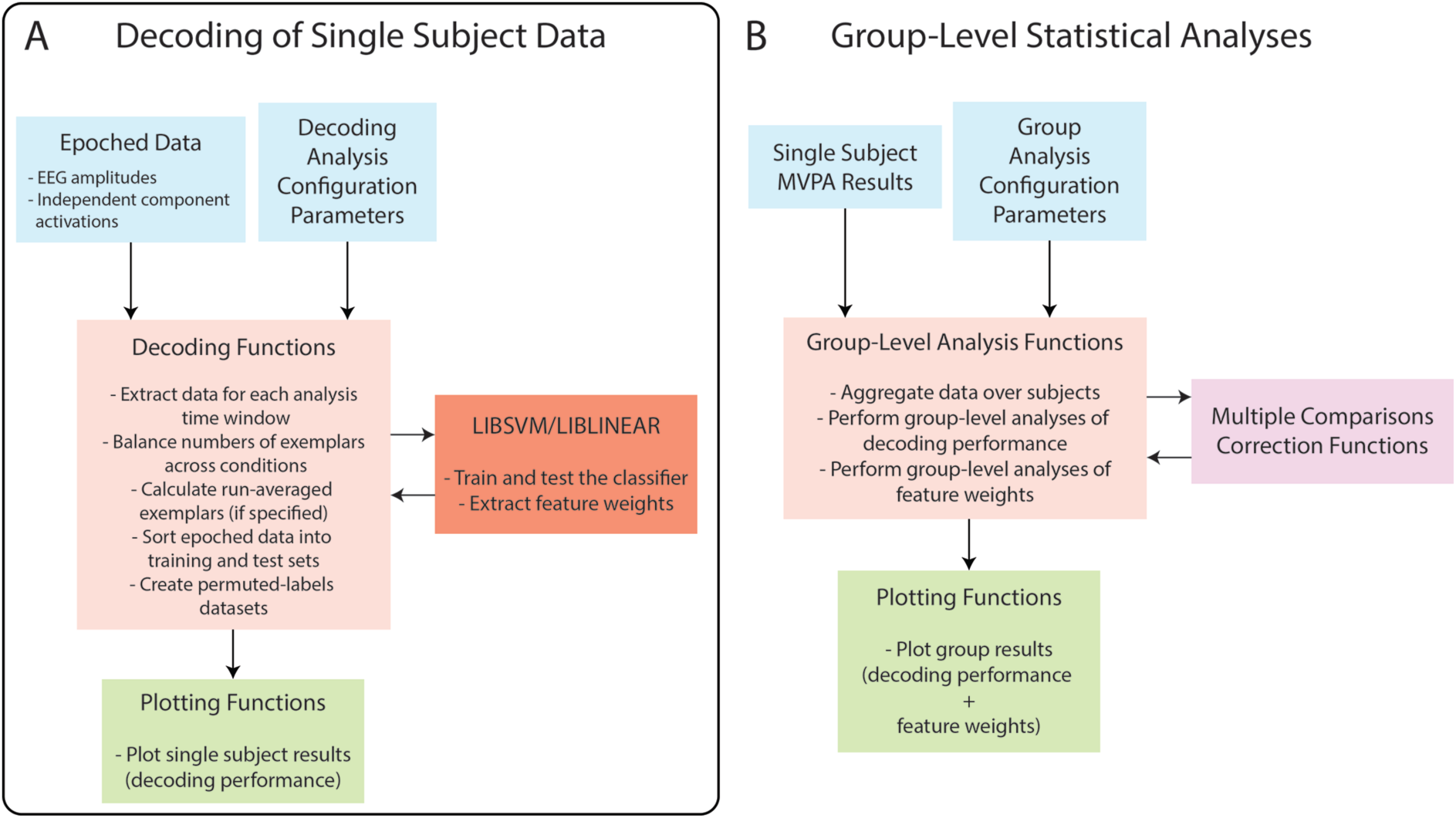
Functional structure of DDTBOX. A) The single subject data decoding functions accept epoched data and analysis configuration parameters. Epoched data is extracted for selected analysis time windows, and sorted for SVM classification or regression, for each cross-validation step and each independent analysis (full set of cross-validation steps). SVM classification/regression is performed in LIBSVM or LIBLINEAR. B) Group-level statistical analysis functions accept single subject MVPA results and group analysis configuration parameters. Decoding performance and feature weights are aggregated over single subjects and are statistically tested at the group level. Multiple comparisons corrections are applied as specified by the user. After analyses, DDTBOX can plot the group decoding accuracy and feature weights results.

The following section will only provide a brief overview of the functional structure, which is divided into phases:

*Data Preparation (Phase 1).* Includes preparation of the epoched data (see Section 5.1 below), as well as configuration of classification/regression analyses (as covered in the previous section).

*Reading the data (Phase 2).* This data is transformed into a MATLAB cell array with the following format:
`eeg_sorted_cond{run, cond}(timepoints, channels, trials)`

whereby *run* refers to the experimental block (if no separate blocks exist in the data, *run* will be 1), *cond* is the category/condition for classification (only one condition is used for support vector regression), *timepoints* are the single data points, *channels* the included EEG channels, and *trials* the single trials of the experiment. This is the general format for data storage, and each processing step will create a similar variable after the specified manipulations.

*Reduction of data (Phase 3).* Next, the data is reduced to the user-specified categories / conditions, which are used for the discrimination group of interest. This has the advantage that DDTBOX can operate within the memory constraints of most desktop computers.

*Balancing the number of included trials (Phase 4).* A frequent problem with classification analysis is that one might end up with an unequal number of trials per condition. This might be due to paradigms in which one condition is over-represented (e.g., oddball paradigms, flanker tasks, or any other paradigm that requires more or less frequent events), responses of interest are not balanced (e.g., errors and correct responses, or most decision-making paradigms), or simply because by chance more trials are lost during EEG data pre-processing for one than for another condition. While this is not necessarily a problem for classification analyses, DDTBOX takes a conservative approach and equalises the number of trials per category / condition before classification.

*Calculating block-average trials or pooling all trials across blocks (Phase 5).* The next step involves averaging across trials (i.e. exemplars) within each experimental block, if this option was chosen. Alternatively, if there exist multiple blocks, but the user chose to treat them all as one long experiment, trials from each block are pooled at this stage.

*Sorting for classification (Phase 6).* The data is now sorted for the classification or regression process. For this, all trials (or block-averaged trials) will be divided into the user-specified number of k sets (the default is k = 10), which also specifies the number of cross-validation sets to be executed. For each full cross-validation cycle (repeated m times; the default is m = 10; see section *3.7 Statistical testing*) trials are randomly assigned to one of the sets with the restriction that no set can have more trials than the others (left-over trials are excluded for this cycle). Of these sets, k – 1 are randomly assigned to the training data variable while the left-out set is assigned to the test data variable. All k combinations are stored before the random assignment of trials to sets and their sorting into training data and test data is performed again for all m iterations. For SVR, an additional matrix containing one value (the condition label) for each trial is used and substitutes for the class labels.

*Vector preparation (Phase 7):* After sorting data into training and test sets, DDTBOX extracts data from within the analysis time window and reshapes data from each trial into a single vector. These vectors are then used for training and testing the SVM classification or regression model.

## 5. Using DDTBOX

Preparing and running MVPA in DDTBOX involves four stages: preparing the data, configuring and running the decoding analyses, configuring and running group-level analyses, and plotting and interpreting the group results. Each of these are briefly described below.

### 5.1 Preparation of EEG data

For decoding analyses DDTBOX uses epoched data, as described in section 4. Each participant dataset is saved as a separate data file. Epoched EEG data must be sorted by experimental condition and run/block, and then stored in this array. If applicable, SVR labels are stored within a separate cell array, with labels ordered in the same way as the corresponding epochs in the EEG data array. A function for automatically converting EEG data epoched using EEGLAB or ERPLAB is provided with the toolbox. This function can also extract epoched independent component activations in addition to EEG amplitudes. This function can further generate SVR labels files for each condition. Other data types (such as behavioural or steady-state visual evoked potential data) can, in principle, also be organised within the same cell array structure for use with DDTBOX by advanced users (for more information see the online documentation).

### 5.2 Configuring and running the decoding analyses

DDTBOX uses a decoding analysis configuration script for defining all relevant parameters and running decoding analyses. Within this script the user can define single subject data filepaths, EEG dataset information, experimental conditions and discrimination groups, and a wide variety of multivariate classification/regression analysis parameters. Finally, the subjects and discrimination groups for analyses are defined, and the DDTBOX core decoding functions are called from this script. Users can copy and adapt these scripts for their own experiments; all parameters are clearly explained in the code comments of the script.

Once all the configuration parameters have been specified, the user can run decoding analyses by executing the MATLAB configuration script. SVM classification/regression performance and feature weights information will be stored in a separate file for each subject.

### 5.3 Configuring and running group-level analyses

Group-level statistical analyses of classification/regression performance and feature weights are configured and run using a group-level analysis configuration script. Within this script the user must define the filepaths of decoding results files, EEG dataset information, group-level statistical analysis and plotting parameters, and must specify the subjects and discrimination groups to use for analyses. Running this configuration script will perform all specified group-level statistical analyses on classification/regression performance and feature weights, which can also be plotted at this stage if desired.

### 5.4 Plotting and interpreting the group results

DDTBOX offers a variety of plotting options for classification/regression performance and feature weights results at the group level. These may be performed when running group-level statistical analyses, and can be replotted using a separate set of easy-to-configure plotting scripts.

For *spatial* and *spatiotemporal decoding analyses* group average classification/regression performance is plotted for each selected time window in the epoch, for results of both original and permuted labels decoding analyses (Figure 3A). For *temporal decoding analyses* group average performance for a single analysis time window is plotted as a topographic heat map (Figure 3B). Feature weights are also plotted in this way, and can also be plotted as a map thresholded for statistical significance (Figure 3C).

**Figure 3.**
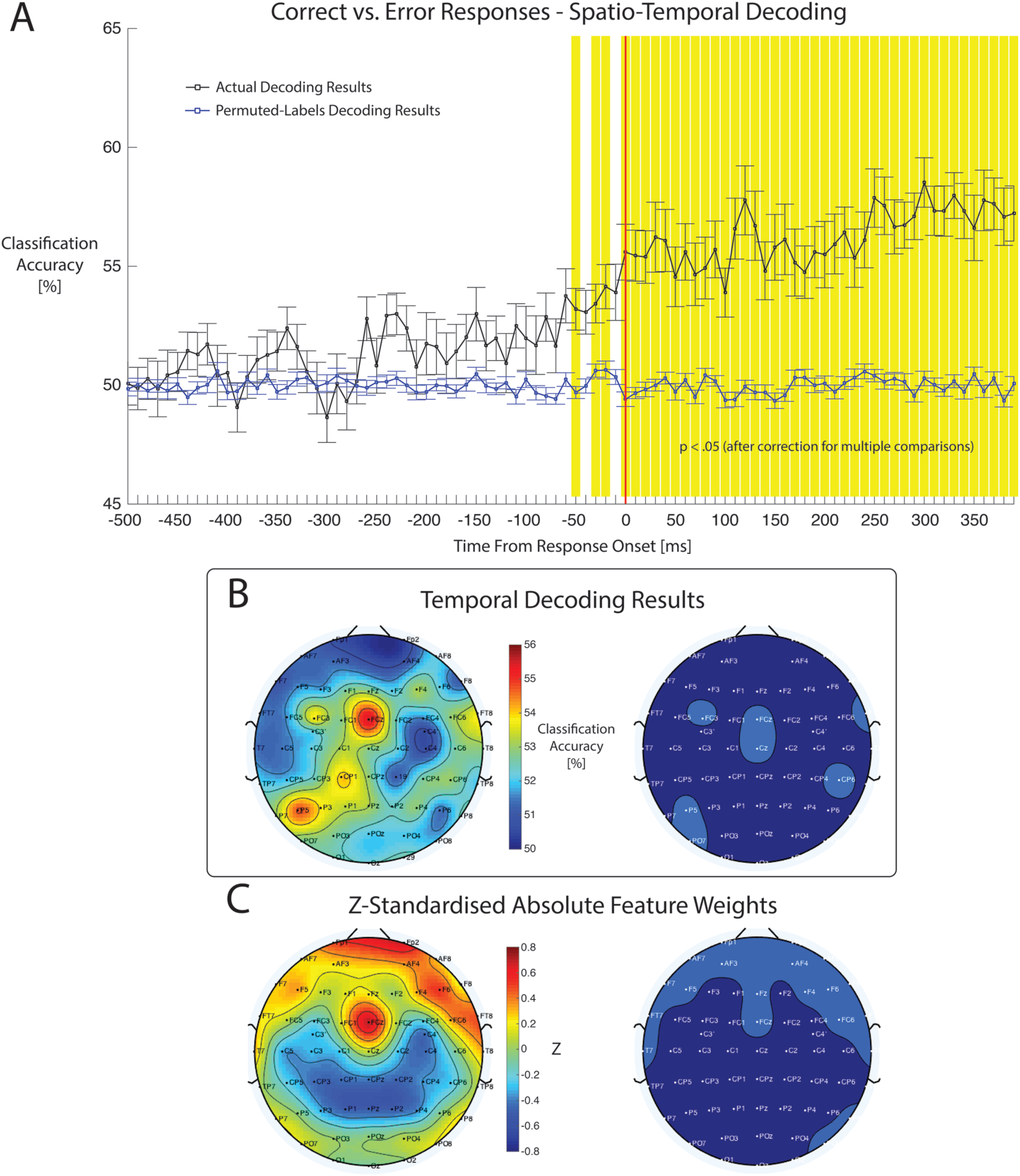
Examples of group-level results outputs produced by DDTBOX. A) Group average classification accuracy scores by time window from response onset. The black line represents the actual decoding results, blue line is the permuted-labels analysis results. Error bars represent standard errors of the mean. Shaded time windows are statistically significant after correction for multiple comparisons. B) Temporal decoding results. A single time window was selected for temporal decoding analyses (100-300ms from response onset). This time range approximates the timing of the error positivity ERP component in Bode and Stahl (2014). The left scalp map plots group average classification accuracy for each channel. The map on the right highlights channels showing decoding accuracy scores that were statistically significantly above zero. C) Feature weights results averaged over time windows spanning 100-300ms from response onset. The left scalp map displays group averages of z-standardised absolute feature weights. The map on the right highlights feature channels with feature weights with z-scores that were significantly above zero.

## 6. Examples of research using DDTBOX

In this section, we briefly review some studies that have used DDTBOX to investigate cognitive functions. We will use these to illustrate some recent research questions for which MVPA analysis has been profitably applied to ERP data; however, there are many other potential research questions for which DDTBOX could be used.

DDTBOX owes its name to its first application in perceptual decision-making (Bode et al., 2012). In this EEG study, images of pianos and chairs were presented after a 100ms forward mask and longer backward mask (500ms minus the duration of the target stimulus, which was either 16.7ms, 33.3ms, 50ms, or 66.7ms). A randomised response mapping screen was shown after the backward mask, circumventing early motor preparation. DDTBOX was used to predict the displayed object category, as well as participants’ category choices, at all four discriminability levels. First, a spatial classification approach was applied, using 80ms analysis time windows moved in steps of 20ms. It was found that the spatial patterns of EEG data predicted the displayed as well as the chosen category during the presentation of the poststimulus mask, with decreasing accuracy and fewer predictive time windows with decreasing discriminability of the objects (Bode et al., 2012). The study also presented phase-randomised visual noise images at the shortest presentation duration (16.7ms), but participants believed themselves to be guessing real object categories. Participants’ choices could be predicted from activity patterns from the pre-stimulus time period. This was interpreted as brain activity reflecting pre-existing decision biases resulting from carry-over effects of decisions in previous trials. To identify channels likely to contain this predictive information, complementary temporal classification analyses, using data from each channel separately, were performed for selected time windows showing high group classification accuracy in the spatial decoding analysis. Temporal decoding analyses showed that channels predominantly over the visual cortex encoded object information early after stimulus presentation, while prefrontal channels did so during later stages before response preparation. For the pure-noise condition, decision-related information was found for both channels over visual cortex and prefrontal cortex during the pre-stimulus period. Taken together, these results demonstrate that the classification analyses as implemented in DDTBOX can indeed detect subtle decision-related information, which would have gone unnoticed in conventional ERP analyses. In contrast to the MVPA results, no ERP components selected for analyses showed differential activity related to piano and chair decisions, or differences in prestimulus baseline activity by category decision for the pure noise condition. This is likely due to subject-specific patterns of EEG activity that differ in response to pianos and chairs, which may not be consistent across subjects and ‘average-out’ in conventional univariate ERP analyses.

In another EEG study, spatiotemporal classification was performed using DDTBOX to predict whether an upcoming response for a parity decision in a speeded digit flanker task was correct or erroneous (Bode and Stahl, 2014). Participants were asked to indicate, using one of two response buttons, whether a central digit on the screen was odd or even, in the presence of two flanker digits on each side that were also odd or even, thereby creating congruent or incongruent decision conditions. For MVPA 10ms analysis time windows were used, moved in steps of 10ms through the trial, approaching the behavioural response onset. MVPA revealed that EEG activity patterns from 100ms before response execution already predicted whether the upcoming response would be erroneous, while conventional ERP analyses found that the error-related negativity (ERN), which follows a response by 80-100ms, was the first ERP component to predict decision errors (Bode and Stahl, 2014). Follow-up analyses of feature weights suggested that this early information originated from channels over visual and motor cortices. In this study classification analyses performed using DDTBOX provided information related to decision errors preceding the participants’ responses, and informed theories of how information about upcoming decision errors could accumulate over time to support online error monitoring processes (Bode and Stahl, 2014).

DDTBOX has also been used to investigate perceptual categorisation of faces (Quek and Rossion, 2017) and multi-sensory integration in elderly and younger adults (Chan et al., 2017). Another application of DDTBOX has been the use of SVR to predict postexperimental ratings of affective and abstract stimulus attributes of task-irrelevant images to inform theories of automatic processing of stimulus features during passive exposure (Bode et al., 2014). These latter examples demonstrate that DDTBOX is by no means restricted to applications in decision-making. On the contrary, it lends itself to many possible questions for which conventional ERP analyses might not be suited, such as cases in which the specific encoding patterns and the timing of these patterns are unknown prior to the experiment.

## 7. Limitations, future developments and extensions

Although it includes a range of analysis options, the current version of DDTBOX is still limited in several ways. The first notable limitation is the support for MVPA using different types of input data. At this stage, DDTBOX can perform MVPA on frequency domain and time-frequency data, as well as component activations from principal components analysis (PCA) or independent components analysis (ICA). However, DDTBOX does not yet offer result plotting capabilities, or automatic conversion to DDTBOX-compatible data files, for these data types. Future support for these data types will widen the applicability of DDTBOX for use with different experimental designs, for example studies examining multivariate patterns of steady-state visual evoked potential (SSVEP) data (e.g., Jacques et al., 2016). In particular, decoding with principal or independent components may also help improve decoding accuracy compared to EEG amplitudes (Grootswagers et al., 2016).

Another current limitation of DDTBOX is its restriction to using the same analysis time window for training and testing. Others have suggested that one strength of the multivariate approach is that the temporal generalisability of patterns across time can be investigated (Meyers et al., 2008; Carlson et al., 2011; King and Dehaene, 2014; Fahrenfort et al., 2017). For this, a classifier could be trained on data from one time window and then tested at other time windows to assess the duration for which the same training data successfully predicts the cognitive process (or content) of interest. By using all possible combinations of training and test data, a full generalisation matrix can be compiled that is informative about the temporal dynamics of cognition (c.f. Fig. 3 in King and Dehaene, 2014; see also Hoogendorn, 2015). Temporal generalisation analyses will be added to a future version of DDTBOX.

A final noteworthy limitation is that the user is required to extract epoched data from EEGLAB/ERPLAB, and to create a configuration script containing all necessary information about the study and planned analyses, before using DDTBOX. While we provide a user-friendly wiki, example configuration scripts, and functions for automatically extracting data epoched using EEGLAB/ERPLAB, the use of DDTBOX nevertheless requires some basic knowledge of MATLAB. Our aim is that the next release will also function as a plug-in for EEGLAB, providing users with a graphical user interface (GUI) within the EEGLAB environment to input all DDTBOX configuration parameters, and the option to use data directly from EEGLAB. However, we are confident that the current release will be of great benefit for the research community, and our toolbox can easily be handled without a GUI.

## 8. Summary

To conclude, DDTBOX is a freely available, open-source toolbox for MATLAB that can be used for multivariate pattern classification and regression analyses on spatial, temporal and spatiotemporal patterns of EEG data. It is useful for investigating cognitive processes related to decision-making, object categorisation, perception, and potentially many other cognitive phenomena. This class of predictive methods can be used in a more explorative and data-driven fashion than conventional ERP analyses. DDTBOX has been used in several published studies and allows for detecting even subtle information that might be overlooked by standard ERP analyses. DDTBOX incorporates a variety of statistical tests, and the option to perform permuted-labels analyses to generate empirical chance distributions. It also generates feature weight maps, which provide useful estimates of the origins of the decodable information.

DDTBOX and the respective documentation is available at: <u>https://github.com/DDTBOX/DDTBOX.</u>

The developers are working on improving DDTBOX on a regular basis. Users can subscribe to our mailing list and will be regularly updated about new releases and features. As the code is openly-available on GitHub, we invite all users to contribute to DDTBOX by submitting their own extensions and improvements. Authors of accepted contributions will be acknowledged in future releases. With DDTBOX, we are hoping to provide a useful toolbox for multivariate EEG analysis that can grow with the needs of researchers and new directions in the field, driven and developed further by an active community of users.

## 9. Author contributions

The DDTBOX has been developed and written by SB, with significant contributions by DB, DF and PMA. All authors contributed to the online documentation and developed the learning material. SB, DF wrote the paper. All authors contributed to and approved the final version of the paper and agreed to be accountable for the content of this work. Please contact for questions regarding DDTBOX.

## 11. Acknowledgements

The DDTBOX was inspired by SB’s work with Prof John-Dylan Haynes on MVPA for fMRI, and some features of the code were modelled from code developed in the Haynes lab. We acknowledge helpful input from Dr Carsten Bogler and Dr Chun Siong Soon during this time. We are further thankful for important conceptual input and improvements resulting from collaborative work with Prof Jutta Stahl, Dr Simon Lilburn, Prof Philip L. Smith, Dr Elaine Corbett, Dr Carsten Murawski and Dr Owen Churches.

### 12. Conflict of interest statement

The authors declare no conflict of interest. No payments were received by neither the institutions nor funding agencies to create this toolbox, and institutions and funding agencies had no input into the content of the work or the publication. This toolbox is licensed under the GNU Public License (GPL) version 2.

